# CHCHD4 regulates a proliferation-EMT switch in tumour cells, through respiratory complex I mediated metabolism

**DOI:** 10.1101/513531

**Authors:** Luke W. Thomas, Cinzia Esposito, Jenna M. Stephen, Ana S. H. Costa, Christian Frezza, Thomas S. Blacker, Gyorgy Szabadkai, Margaret Ashcroft

## Abstract

**BACKGROUND:** Mitochondrial metabolism involves oxidative phosphorylation (OXPHOS) via the respiratory chain and is required for the maintenance of tumour cell proliferation and regulation of epithelial–mesenchymal transition (EMT)-related phenotypes through mechanisms that are not fully understood. The essential mitochondrial import protein coiled-coil helix coiled-coil helix domain-containing protein 4 (CHCHD4) controls respiratory chain complex activity and oxygen consumption, and regulates the growth of tumours *in vivo*. In this study we interrogate the role of CHCHD4-regulated respiratory chain activity and metabolism in tumour cell proliferation and EMT-related phenotypes.

**RESULTS:** We show that *CHCHD4* is essential for the proliferation of tumour cells irrespective of their aetiology. In human tumours, elevated *CHCHD4* expression is correlated with a mitochondrial OXPHOS gene signature and with a proliferative gene signature associated with the mTORC1 signalling pathway. Elevated CHCHD4 increases tumour cell proliferation, in a manner that is dependent on complex I (CI) activity, glutamine consumption and mTORC1 activation. CHCHD4 expression is inversely correlated with EMT gene expression both in vitro and in vivo. Finally, we show CHCHD4 regulates the intracellular distribution of the EMT marker vimentin, in a CI-mediated manner.

**CONCLUSIONS:** CHCHD4 regulates tumour cell proliferation and metastatic (EMT-related) phenotypes through its control of CI-mediated mitochondrial metabolism.

## INTRODUCTION

Rapidly dividing tumour cells require specific metabolites to support proliferation, and the metabolic rewiring of malignant cells contributes both to transformation and tumour progression [1]. One of the earliest observations of metabolic adaptation in tumour cells came from the work of Otto Warburg, who identified that even in the presence of sufficient oxygen, many cancer cells consumed high concentrations of glucose and secreted high levels of lactate [2]. In addition to oncogene-driven increases in glucose consumption, tumour cells also increase their consumption of glutamine [3], as glutamine provides carbon and nitrogen moieties for amino acid and nucleotide synthesis. Despite the prevalence of oxidative fermentation in transformed cells, tumour cells retain their oxidative mitochondrial machinery to support the catabolism of glucose and glutamine for the production of macromolecules to permit cell division [4, 5]. Indeed, a variety of recent studies have demonstrated that while the availability of adenosine triphosphate (ATP) is unlikely to be a limiting factor for the proliferation of tumour cells [6–8], the availability of amino acids and nucleotides can be, depending on the cellular context [8–12]. However, our understanding of the mechanisms by which transformed cells maintain biosynthesis of macromolecules for cell division remain incompletely understood. Delineating the pathways which support tumour cell proliferation are of great interest for the development of new therapeutic strategies, and anti-tumour agents which inhibit nucleotide synthesis (e.g. Fluorouracil) have been used clinically for decades [13]. Therapies which target amino acid synthesis are emerging in clinical development, and are showing promise [14–16].

Mitochondria support cellular proliferation by supplying ATP for the bioenergetic demands of the cell through OXPHOS, and are also the site of reactions which supply the cell with precursors for the synthesis of macromolecules such as DNA, proteins and lipids [17]. In addition, complex (C)I regulates the intracellular NADH/NAD ratio, which is itself an essential cofactor for biosynthetic reactions which support proliferation [18]. Depletion of mitochondrial DNA in tumour cells (ρ^0^ cells) inhibits tumour cell proliferation and tumorigenesis in vitro and in vivo, demonstrating the importance of mitochondria in a cancer setting [19–21]. Furthermore, CI-inhibiting biguanidines (e.g. metformin) are effective in slowing the growth of tumour cells in vitro [22, 23], and are under investigation in clinical trials as adjuvant therapies for cancer patients.

All but 13 subunits of the respiratory chain complexes are encoded by nuclear genes, and must be imported across the mitochondrial membrane(s) for complex assembly to take place. Several import and sorting pathways exist in the mitochondria which are essential for respiratory chain activity [24]. One such pathway, the disulphide relay system (DRS) is responsible for the import and oxidative folding of small intermembrane space (IMS) proteins, which include subunits of CI and CIV, as well as assembly factors for CIII and CIV [25–27]. The substrate-binding oxidoreductase of the DRS is the protein CHCHD4, which we have previously shown to regulate oxygen consumption rates, and hypoxia responses through effects on HIF signalling [28, 29]. We have also shown that shRNA-mediated silencing of *CHCHD4* reduces the expression of respiratory chain subunits (CI-CIV) in tumour cells [30] and significantly decreases tumour growth in vivo [28]. In addition, we have shown that elevated *CHCHD4* expression correlates with increased tumour grade and decreased survival of breast cancer patients, demonstrating an association with disease progression [28]. In this study we identify mechanisms by which CHCHD4 controls tumour cell proliferation through its effects on mitochondrial metabolism.

## RESULTS

### CHCHD4 expression correlates with mitochondrial OXPHOS and proliferative gene signatures in vitro and in vivo

*CHCHD4* is an essential gene in mice [31]and yeast (*Mia40*) [32], and here, we have found that *CHCHD4* is ranked as an essential gene for the proliferation of a panel of 341 tumour cell lines from the Broad Institute Achilles Project (Fig. 1a). To better understand the relationship between CHCHD4 and tumour cell proliferation, we carried out gene set enrichment analysis (GSEA) on genes that were significantly correlated with *CHCHD4* expression in transcriptomic data from a panel of 967 tumour cell lines (Novartis/Broad Institute Tumour Cell Line Encyclopaedia). As we anticipated, *CHCHD4* was significantly co-expressed with genes from an OXPHOS gene set (HALLMARK_OXIDATIVE_PHOSPHORYLATION, Broad Institute) (Fig. 1b), including subunits of CI (*NDUFS3*) and CIV (*COX7C*) (Additional File 1: Figure S1a). Interestingly, many of the most significantly enriched gene sets correlated with *CHCHD4* were those from proliferative signalling pathways regulated by known oncogenes, including MYC and mTORC1 (Fig. 1b). Genes regulated by mTORC1 signalling that were most significantly correlated with *CHCHD4* expression included cell-cycle regulators (*CDC25A, CCNF*) and subunits of DNA polymerases (*POLR3G*) (Additional File 1: Figure S1b). We next carried out GSEA on genes that were significantly co-expressed with *CHCHD4* in different cancer cohorts, from transcriptomic data available from The Cancer Genome Atlas. Again, we found that *CHCHD4* was significantly co-expressed with genes involved in OXPHOS in colon adenocarcinoma (Fig. 1c), breast cancer (Fig. 1d) and glioblastoma (Additional File 1: Figure S1c) patient samples. In colon adenocarcinoma (Fig. 1e) and breast cancer patient samples (Fig. 1f), the most highly correlated genes involved in OXPHOS constituted components of respiratory chain complexes (e.g. *UQCRH*, *NDUFAB1*), as well as enzymes of the TCA cycle (*DLAT*, *FH*), and proteins involved in the general maintenance of mitochondrial function (e.g. *TOMM70*, *HSPA9*). *CHCHD4* expression was positively correlated with the expression of genes regulated by proliferative signalling pathways such as MYC and mTORC1 in each of the patient cohorts analysed (Fig. 1c, d, Additional File 1: Figure S1c). Genes regulated by mTORC1 signalling that were co-expressed with *CHCHD4* in colon adenocarcinoma patient samples included cell cycle regulators (*CCNA2, CDC25A*), and genes involved in nucleotide synthesis (*MTHFD2L*), suggesting a potential relationship between *CHCHD4* expression and mTORC1 regulation of proliferation (Fig. 1g).

**Figure 1.**
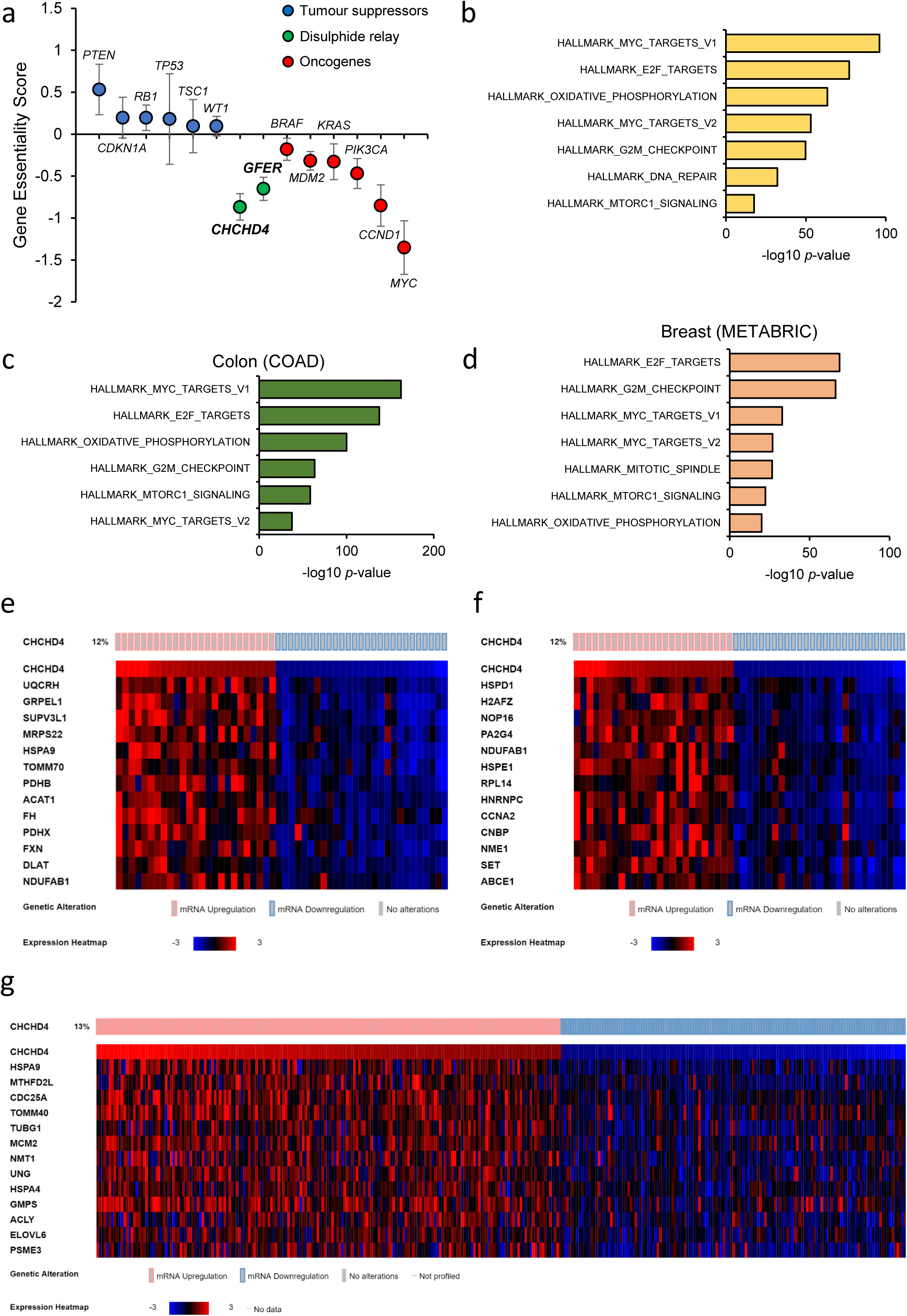
CHCHD4 expression correlates with OXPHOS and proliferative gene signatures in vitro and in vivo. **a** Gene essentiality scores derived from Broad Institute Achilles Project for selected tumour suppressors, oncogenes, and *CHCHD4* and *GFER*. ±SD. n=341 cell lines. **b** Chart shows GSEA of genes positively correlated with *CHCHD4* expression in Novartis/Broad Institute Cancer Cell Line Encyclopedia RNASeq data. n=967 cell lines. **c-d** Chart shows GSEA of genes positively correlated with *CHCHD4* expression in colon adenocarcinoma (**b**) and breast cancer (**c**) patient tumours. **e** Heatmap of selected genes from HALLMARK_OXIDATIVE_PHOSPHORYLATION gene-set (Broad Institute) that are positively correlated with *CHCHD4* expression in colon adenocarcinoma patient tumours. **f-g** Heatmap of selected genes from HALLMARK_MTORC1_SIGNALLING gene set (Broad Institute) that are positively correlated with *CHCHD4* expression in colon adenocarcinoma (**f**) and breast cancer (**g**) patient tumours.

### Tumour cell proliferation correlates with CHCHD4 expression and mitochondrial function

Mitochondrial function is essential for the maintenance of biosynthetic pathways and our GSEA analysis suggested that proliferative signalling pathways were correlated with CHCHD4 expression and the expression of genes regulating OXPHOS, both in vitro and in vivo (Fig. 1). To further investigate the relationship between CHCHD4, OXPHOS and tumour cell proliferation, first we tested a panel of cell lines to identify tumour cells that exhibited differing dependencies on OXPHOS for proliferation. To do this, we selected six well known tumour cell lines derived from different tissues (colon, cervix, bone, breast, brain and prostate), and each with a different oncogenic background. We initially assessed their relative rates of proliferation (Fig. 2a) and ranked the cell lines by proliferative capacity. Next, to characterise the dependence of these cell lines on mitochondrial OXPHOS for their proliferation, we assessed the relative sensitivity of these cell lines to respiratory chain inhibition using titrations of the CI inhibitors rotenone (Fig. 2c) and BAY 87 2243 [33] (Additional File 2: Figure S2a), and the CIII inhibitor antimycin A (Additional File 2: Figure S2b). Sensitivity to these agents was inversely correlated with the relative rates of proliferation of these cell lines (e.g. rotenone, Fig. 2c), demonstrating the common importance of respiratory chain activity for tumour cell line proliferation. We next assessed CHCHD4 levels in two cell lines from our panel, one with a high proliferation rate and increased sensitivity to respiratory chain inhibition (HCT116 colon carcinoma), and one with a lower proliferation rate and reduced sensitivity to respiratory chain inhibition (U2OS osteosarcoma). Interestingly, CHCHD4 expression was higher in HCT116 cells compared to U2OS cells both at the transcript level (Fig. 2d) and protein level (Fig. 2e). Furthermore, the expression of subunits of each of the respiratory chain complexes (CI-IV) was higher in HCT116 cells compared to U2OS cells (Fig. 2e), as was basal oxygen consumption rate (Fig. 2f) demonstrating increased mitochondrial function. Collectively, our data support a relationship between CHCHD4 expression, mitochondrial OXPHOS and proliferative capacity irrespective of tumour cell context, as also demonstrated in our transcriptomic analyses (Fig. 1).

**Figure 2.**
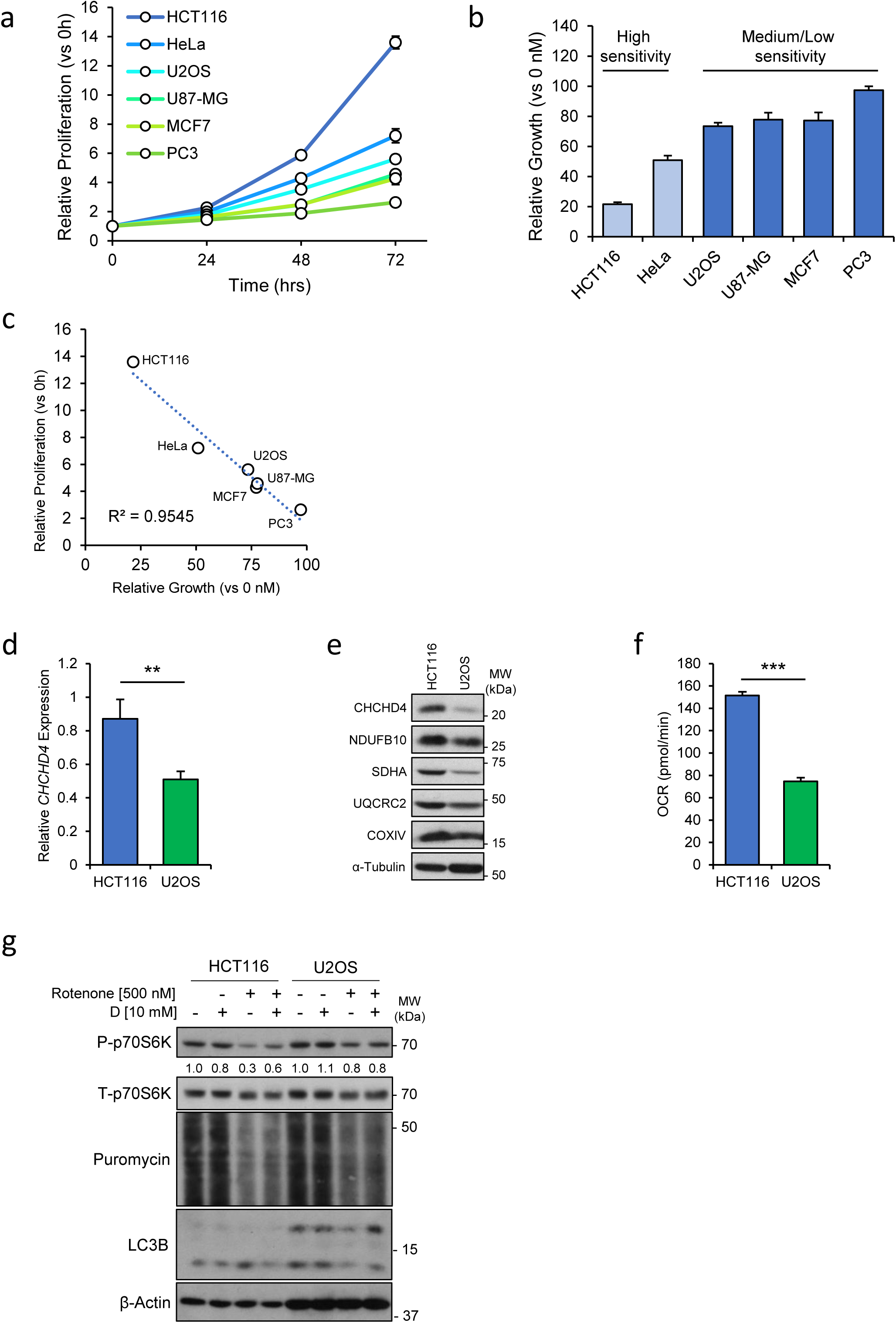
Tumour cell proliferation correlates with CHCHD4 expression and mitochondrial function. **a** Chart shows proliferation rates of tumour cell line panel over 72h. ±SD. n=3. **b** Chart shows growth of tumour cell line panel treated with 100 nM rotenone for 72h, relative to untreated (0 nM) cells. ±SD. n=3. **c** Chart shows xy scatter of proliferation rates of indicated cell lines at 72h, and relative growth rates treated with 100 nM rotenone. Trend line (dashed blue) and R^2^ value (Spearman’s correlation) shown. **d** Chart shows relative abundance of *CHCHD4* transcripts measured by QPCR in HCT116 cells and U2OS cells, relative to *ACTB* and normalised to HCT116 transcript levels. ±SD. n=3. **e** Western blot shows levels of CHCHD4, NDUFB10 (CI), SDHA (CII) UQCRC2 (CIII), COXIV (CIV) in whole cell lysates of HCT116 and U2OS cells. α-Tubulin used as load control. **f** Chart shows basal OCR of HCT116 and U2OS cells. ±SD. n=3. **g** Western blot shows levels of phosphorylated (P-) and total (T-) p70S6K, puromycin labelled polypeptides, and LC3B in HCT116 and U2OS cells treated as indicated for 24h. β-Actin used as load control. Relative band intensities of P-p70S6K indicated.

Mitochondrial respiration supports proliferation through the production of ATP, and also through production of NAD (at CI), which is an essential cofactor for amino acid and nucleotide biosynthesis [18]. Nutrient availability is co-ordinated with proliferation through several signalling pathways, including mTORC1 signalling [34], which we found was one of the most significantly enriched pathways correlated with *CHCHD4* expression in vitro and in vivo (Fig. 1). In nutrient sufficient conditions, mTORC1 is activated and promotes protein translation through phosphorylation of downstream targets including p70S6 Kinase 1 (p70S6K), as well as inhibition of autophagy [34]. We next investigated the contribution of the mTORC1 pathway to mitochondrial OXPHOS inhibition using HCT116 and U2OS cells from our cell line panel and the CI inhibitor rotenone. We found that rotenone treatment led to a larger decrease in phosphorylation of p70S6K, a larger reduction in de novo protein synthesis (assessed by puromycin incorporation), and accumulation of the lipidated form of LC3B autophagy factor in HCT116 cells compared to U2OS cells (Fig. 2g), indicating a larger reduction in mTORC1 activity in HCT116 cells upon CI inhibition. Through the production of NAD, CI promotes the synthesis of aspartate and asparagine which directly activates mTORC1 [35], and which are also required for the de novo synthesis of nucleotides and other amino acids. Supplementation of these cells with aspartate (D) was capable of partially rescuing the reduced mTORC1 activity in the presence of rotenone (Fig. 2g). Together, our data support a mechanistic relationship between CHCHD4 expression, CI activity, mTORC1 signalling and proliferative capacity irrespective of tumour cell aetiology.

### CHCHD4 regulates CI expression and activity in tumour cells, and regulates the mTORC1 pathway

CHCHD4 is a mitochondrial protein localised to the intermembrane space (IMS), and functions as the substrate-binding component of the DRS, which binds and through its oxidoreductase activity folds subunits and assembly factors for CI, III and IV [27]. In order to further investigate the relationship between CHCHD4 expression and CI-mediated regulation of tumour cell proliferation, we used U2OS cells stably overexpressing wild-type CHCHD4 [CHCHD4 (WT)-expressing cells, clones WT.cl1, WT.cl3], or a mutant form of CHCHD4 in which the substrate-binding cysteines of the CPC motif have been mutated to alanines (C66A/C68A) [28, 29], (Fig. 3a). Western blot analysis of these cell lines demonstrated that wild-type CHCHD4 overexpression increased the expression of individual subunits of each of the respiratory CI-IV (Fig. 3a). Importantly, overexpression of the C66A/C68A mutant form of CHCHD4, which we and others have shown is defective in import function and mitochondrial localisation [28, 36], did not lead to increased respiratory chain subunit expression (Fig. 3a). Interestingly in some experiments we found this mutant markedly decreased the mitochondrial expression of these complex subunits (Fig. 3a), demonstrating that the effects of CHCHD4 on respiratory chain subunit expression is depend on its import function, and also suggesting that this C66A/C68A mutant form of CHCHD4 may function as a dominant negative. To gain a broader understanding of the influence of CHCHD4 on the expression of respiratory chain subunits and the proteome as a whole, we carried out SILAC analyses comparing control U2OS cells with our CHCHD4 (WT)-expressing cells. Our analyses showed that CHCHD4 expression significantly affects CI biology, as we found increased mitochondrial expression of twenty-seven subunits of CI in CHCHD4(WT)-expressing cells compared to control U2OS cells (Fig. 3b), corroborating our western data (Fig. 3a). Furthermore, using an in-gel nitrotetrazolium blue assay, we found that elevated CHCHD4 expression in U2OS cells led to increased activity of whole CI (Fig. 3c), demonstrating that increased CI subunit expression in these cells leads to increased whole CI activity. Notably, in U2OS (C66A/C68A) cells there was a reduction in CI activity (Fig. 3c). Using live cell imaging of NADH fluorescence [37], we next demonstrated that CHCHD4 (WT)-expressing cells had lower basal NADH fluorescence (Fig. 3d), further indicating higher CI activity in these cells. As our data show that sensitivity to CI was correlated with CHCHD4 expression and proliferative capacity (Fig 2), we next assessed the relative effects of CI inhibition (using rotenone) on the proliferation of our control and CHCHD4 (WT)-expressing U2OS cells. Consistent with our data in Fig. 2, we found that elevated CHCHD4 expression in tumour cells led to increased sensitivity to CI inhibitor treatment (Fig. 3e). Conversely, CHCHD4 (C66A/C68A) mutant cells showed similar sensitivity to rotenone as control U2OS cells, indicating that CHCHD4 expression and import function is important for the relative sensitivity of tumour cells to CI inhibitors (Fig. 3e). Consistently, shRNA-mediated silencing of *CHCHD4* decreased the sensitivity of HCT116 cells to growth inhibition by rotenone (Additional File 3: Figure S3a). To further investigate the relationship between CHCHD4, mitochondrial OXPHOS (CI activity), proliferation and mTORC1 signalling, we next carried out GSEA on proteins significantly upregulated CHCHD4 (WT)-expressing cells compared to control U2OS cells from our SILAC analysis. Similar to our findings from patient samples here (Fig. 1), and previously [28], we found that elevated CHCHD4 expression led to a significant increase in the expression of mitochondrial OXPHOS proteins and proteins involved in hypoxia signalling (Fig. 3f). Importantly the most significantly enriched class of proteins upregulated by elevated CHCHD4 were those regulated by the mTORC1 signalling pathway (e.g. GPI, E2F1) (Fig. 3f), again reflecting what we found in cancer patient samples (Fig. 1). To further explore the influence of CHCHD4 on CI biology and mTORC1 activity, we assessed mTORC1 activity in the absence and presence of rotenone treatment in CHCHD4 (WT)-expressing and control U2OS cells. Basal p70S6K phosphorylation and puromycin incorporation were higher in cells overexpressing CHCHD4 (Fig. 3g), indicating higher mTORC1 activity in these cells. Furthermore, rotenone treatment led to a larger reduction in mTORC1 activity in cells overexpressing CHCHD4 (Fig. 3g), and both the increased inhibition of growth (Fig. 3h) and mTORC1 activity (Fig. 3g) could be partially mitigated by supplementation with 10 mM aspartate. Together these data suggest that CHCHD4 regulates tumour cell proliferation, through its effects on CI-dependent activation of mTORC1 signalling.

**Figure 3.**
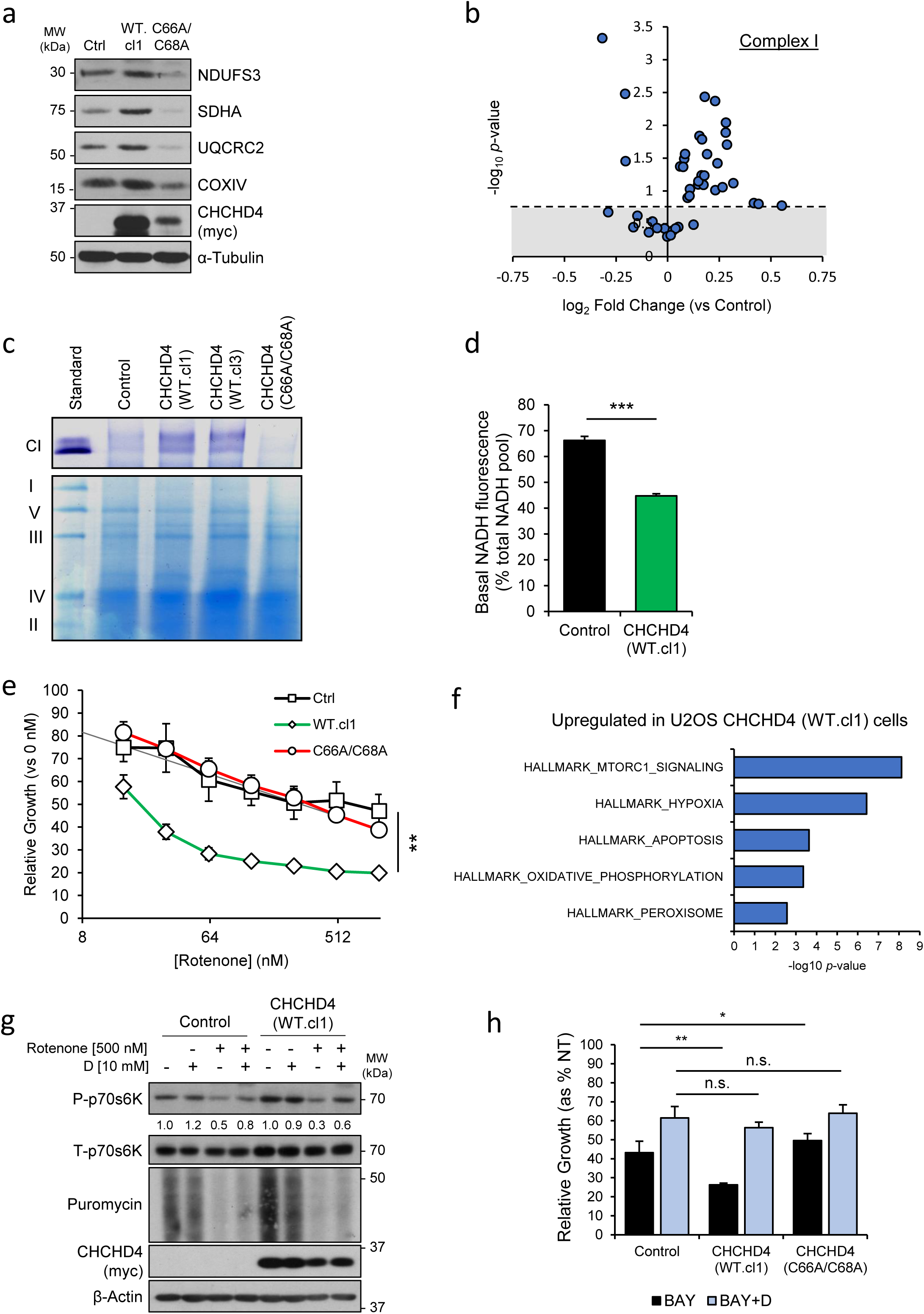
CHCHD4 regulates CI expression and activity in tumour cells, and regulates the mTORC1 pathway. **a** Western blot shows levels of CHCHD4, NDUFB10 (CI), SDHA (CII) UQCRC2 (CIII), COXIV (CIV) in whole cell lysates of control (Ctrl) U2OS cells, and cells overexpressing myc-epitope tagged wild-type (WT.cl1) or mutant (C66A/C68A) CHCHD4. **b** Volcano plot of all detected CI subunit expression in WT.cl1 cells compared to control U2OS cells from SILAC experiments. ±SD. n=3. **c** In-gel NTB assay of CI activity in mitochondrial fractions isolated from control U2OS cells, and cells overexpressing wild-type (WT.cl1, WT.cl3) or mutant (C66A/C68A) CHCHD4. **d** Basal NADH fluorescence (as % of total NADH/NAD pool) in control U2OS cells, and cells overexpressing wild-type CHCHD4 (WT.cl1). ±SD. n=3. **e** Chart shows growth rates of cells described in **a**, treated with a 2-log titration curve of rotenone (starting dose 1µM) for 72h, relative to untreated (0 nM). ±SD. n=3. **f** Chart shows GSEA of proteins upregulated in CHCHD4 (WT.cl1) cells relative to control U2OS cells, as assessed by SILAC. n=3. **g** Western blot shows levels of phosphorylated (P-) and total (T-) p70S6K, and puromycin labelled proteins in control U2OS cells and wild-type (WT.cl1) overexpressing U2OS cells treated as indicated for 24h. β-Actin used as load control. **h** Chart shows relative growth of control U2OS cells, and cells overexpressing wild-type (WT.cl1) or mutant (C66A/C68A) CHCHD4, treated with 10 nM BAY 87 2243 in the absence or presence of 10 mM aspartate (D) for 72h. ±SD. n=3.

### CHCHD4 regulates tumour cell proliferation and glutamine consumption

Based on our data thus far (Fig. 1–3), we hypothesised that CHCHD4-mediated regulation of CI activity stimulates tumour cell proliferation through the promotion of biosynthetic pathways which depend on favourable NAD/NADH ratios, such as the metabolism of glutamine (to amino acids such as aspartate and asparagine), and subsequently leads to the activation of mTORC1 signalling (Fig. 4a). Indeed, the sensitivity of our cell line panel to growth inhibition by glutamine withdrawal directly correlated with their proliferation rates (Additional File 4: Fig. S4a-b), and sensitivity to CI inhibitor treatment (Additional File 4: Fig. S4c). We therefore assessed the influence of CHCHD4 on glutamine-dependent proliferation, by culturing control U2OS cells and CHCHD4 (WT)-expressing cells in the presence and absence of glutamine. Cells overexpressing CHCHD4 had higher proliferation rates than control U2OS cells, and glutamine withdrawal abolished this proliferative advantage (Fig. 4b), suggesting the increased rates of proliferation of cells overexpressing CHCHD4 relied on the availability of glutamine. Furthermore, compared to control U2OS cells, CHCHD4 (WT)-expressing cells were more significantly lost from culture when glutamine was withdrawn (Additional File 4: Fig. S4d), suggesting that tumour cells with elevated CHCHD4 expression depleted their intracellular glutamine pools more rapidly. We next interrogated the effects of CHCHD4 expression on glutamine metabolism by culturing control and CHCHD4 (WT)-expressing cells with uniformly-labelled ^13^C5-glutamine, and analysed both the isotopologue composition and total pool sizes of key glutamine-derived metabolites. We found that intracellular levels of glutamine and other glutamine-derived metabolites were lower in CHCHD4 (WT)-expressing cells compared to control U2OS cells, suggesting greater rates of glutamine consumption (Fig. 4c). However, there were only very minor differences in the isotopologue composition of a selection of these metabolites (Fig. 4d), indicating that there was no significant change in the route of glutamine consumption but only in the rate of glutamine consumption in tumour cells overexpressing CHCHD4. We next assessed the relative influence of glutamine withdrawal on the activity of the mTORC1 pathway in CHCHD4 (WT)-expressing cells, by assessing phosphorylation of p70S6K, and de novo protein synthesis (via puromycin incorporation). Withdrawal of glutamine led to a larger decrease in both p70S6K phosphorylation and puromycin incorporation in cells overexpressing CHCHD4 (Fig. 4e), indicating that the increased mTORC1 activity in CHCHD4 overexpressing cells was dependent on the availability of glutamine. The reduction in mTORC1 activity could be partially reversed by supplementation of the growth media with 10 mM aspartate (Fig. 4e). Similarly, withdrawal of glutamine led to a bigger decrease in mTORC1 signalling in HCT116 cells than in U2OS cells, which was partially recoverable by aspartate supplementation (Additional File 4: Figure S4e). Together these data demonstrate that CHCHD4 regulates the proliferation of tumour cells through increased glutamine consumption and mTORC1 signalling.

**Figure 4.**
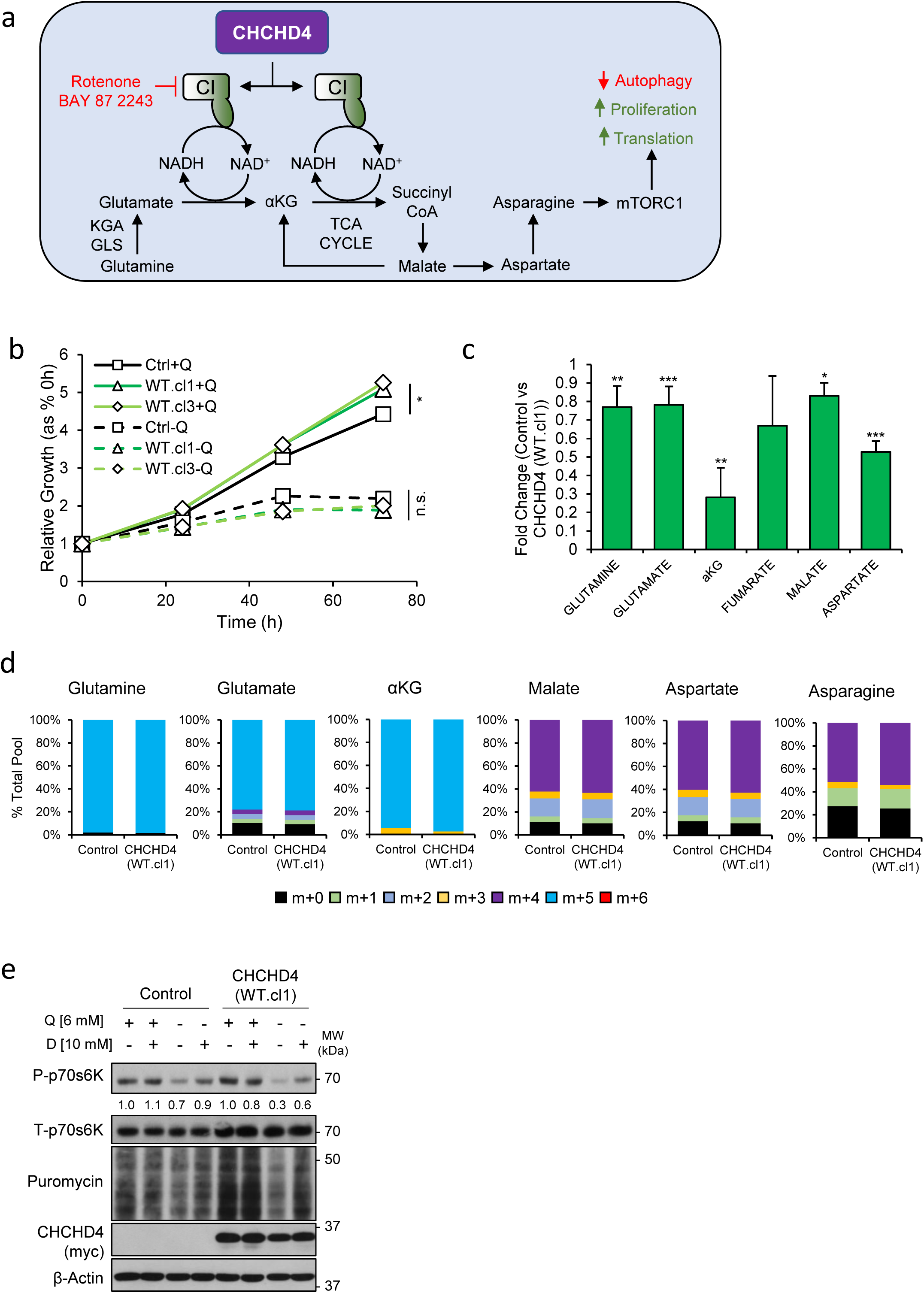
CHCHD4 regulates proliferation and glutamine consumption. **a** Schematic of proposed model of CHCHD4-regulated CI-dependent glutamine metabolism and influence on tumour cell phenotypes, and targets of agents used in this study. **b** Chart shows relative growth rates of control (Ctrl) U2OS cells and U2OS cells overexpressing wild-type (WT.cl1, WT.cl3) CHCHD4, in the presence (+Q) and absence (-Q) of glutamine. ±SD. n=3. **c** Chart shows relative intracellular pool sizes of selected metabolites from metabolomics analysis in WT.cl1 cells relative to control U2OS cells. Representative of 2 incubations. ±SD. n=5. **d** Chart shows relative proportions of isotopically labelled metabolites in control U2OS cells, and WT.cl1 cells. Representative of 2 incubations. n=5. **e** Western blot shows levels of phosphorylated (P-) and total (T-) p70S6K, and puromycin labelled polypeptides in control U2OS cells and wild-type (WT.cl1) overexpressing U2OS cells treated as indicated for 8h. β-Actin used as load control.

### CHCHD4 regulates the EMT phenotype of tumour cells

Along with identifying significantly upregulated genes profiles associated to increased *CHCHD4* expression (Fig. 1), interestingly, our GSEA also showed that epithelial-mesenchymal transition (EMT)-related genes were amongst the most significantly downregulated genes associated with increased *CHCHD4* expression in both patient samples (Fig. 5a, Additional File 5: Figure S5a) and tumour cell lines (Additional File 5: Figure S5b). Recent studies have connected mitochondrial metabolism and health to the EMT phenotype of cultured cells, through changes in cytoskeletal organisation, adhesion and motility [38]. For example, in the context of mitochondrial disease, loss of FH (fumarate hydratase) activity leads to the accumulation of the TCA metabolite fumarate, which competitively inhibits demethylase enzymes, some of which are epigenetic modifiers that control the expression of the antimetastatic *mir-200ba429* miRNA cluster [39]. Indeed, it has long been appreciated that cell proliferation and motility are inversely related, both in physiology and disease [40–42]. We hypothesised that since elevated CHCHD4 increases mitochondrial activity and promotes proliferation, increased CHCHD4 expression might negatively regulate EMT-related genes. Our GSEA of proteins downregulated in cells overexpressing CHCHD4 from our SILAC data identified that EMT-related proteins were the most highly significantly enriched group of proteins (Fig. 5b). These included the essential cytoskeletal protein vimentin, and the extracellular matrix (ECM) binding protein N-cadherin (CDH2), both of which are well-characterised markers of EMT (Fig. 5c) [43, 44]. Furthermore, the intermediate cytoskeleton filament and MET marker keratin-18 (KRT18, [45, 46]) was the second most upregulated protein identified in our SILAC analysis (Fig. 5c). We confirmed some of these EMT protein expression changes in CHCHD4 (WT)-expressing cells by western blot (Fig. 5d, e), while these proteins were unchanged in CHCHD4 (C66A/C68A) expressing U2OS compared to control U2OS cells. We found similar results in HCT116 cells (which do not express detectable levels of vimentin [47]), where overexpression or silencing of CHCHD4 increased or decreased the MET marker E-cadherin expression respectively (Additional File 5: Figure S5c-d). Importantly, we found that the transcript levels of *VIM* and *KRT18* were downregulated and upregulated respectively in cells overexpressing CHCHD4 (Fig. 5f), demonstrating that the changes in EMT gene expression are at the level of transcription, and suggest that elevated CHCHD4 expression in tumour cells leads to a transcriptional suppression of EMT.

**Figure 5.**
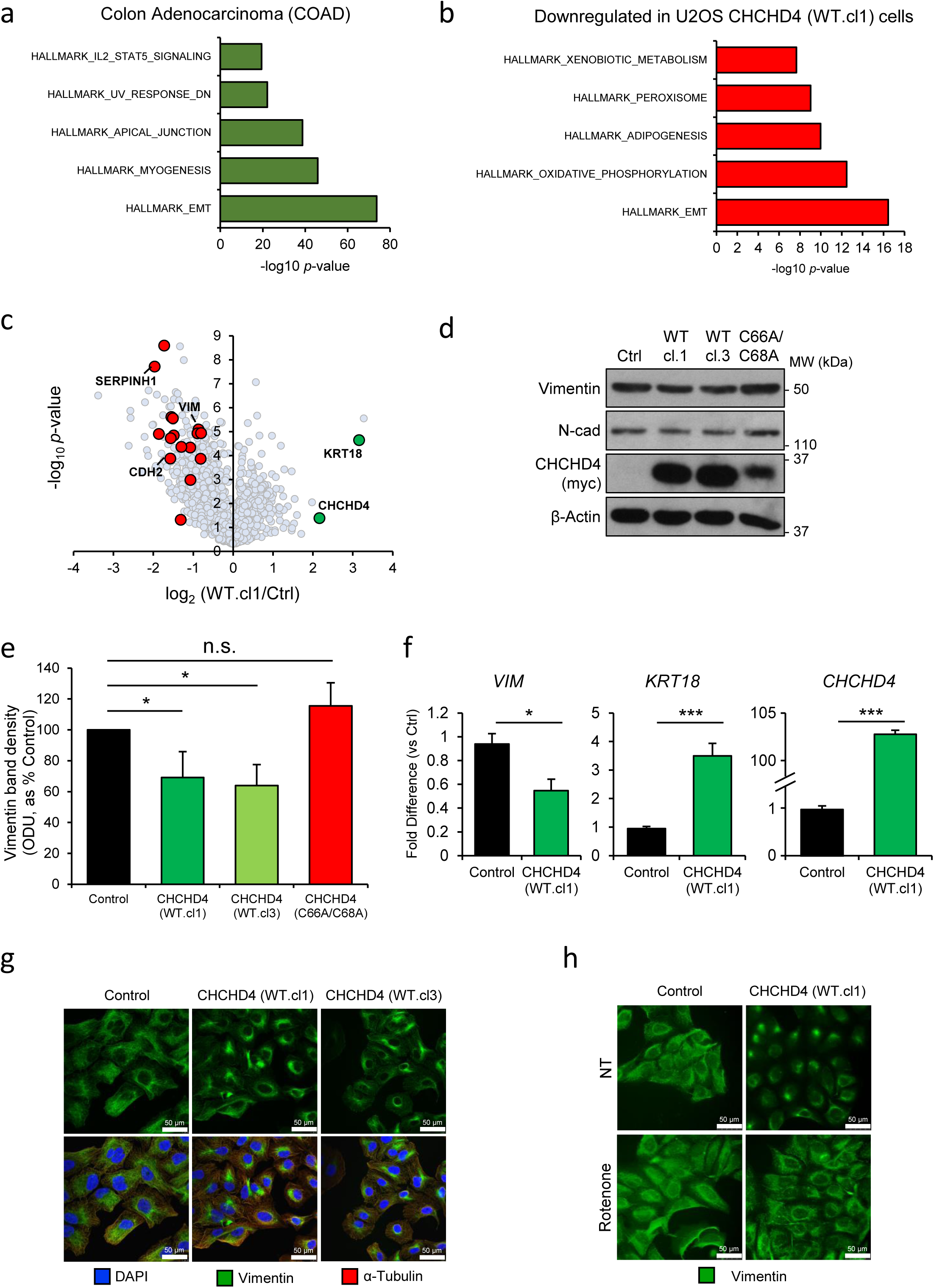
CHCHD4 regulates the EMT phenotype of tumour cells. **a** Chart shows GSEA of genes negatively correlated with *CHCHD4* expression in colon adenocarcinoma patient tumours. **b** Chart shows GSEA of downregulated proteins in WT.cl1 U2OS cells compared to control U2OS cells as assessed by SILAC. n=3. **c** Volcano plot shows relative expression of all proteins detected in SILAC experiments in WT.cl1 cells compared to control (Ctrl) U2OS cells. Selected genes highlighted. n=3. **d** Western blot shows levels of vimentin and N-cadherin in control (Ctrl) U2OS cells, and U2OS cells overexpressing myc-epitope tagged wild-type (WT.cl1, WT.cl3) or mutant (C66A/C68A) CHCHD4. β-Actin used as load control. **e** Chart shows densitometry analysis of vimentin band intensity from 3 independent western blots as described in **d**. ±SD. n=3. **f.** Chart shows relative transcript levels of *VIM*, *KRT18*, and *CHCHD4* detected by QPCR in control and WT.cl1 U2OS cells. ±SD. n=3. **g** Immunofluorescence images of vimentin distribution in control U2OS cells and cells overexpressing wild-type (WT.cl1, WT.cl3) CHCHD4. **h** Immunofluorescence images of vimentin distribution in control U2OS and WT.cl1 cells untreated (NT) or treated with 50 nM rotenone for 72h.

Vimentin is an intermediate filament protein which regulates cell motility through association with other cytoskeletal proteins to promote pseudopodia formation at the cell periphery [48]. Immunofluorescent staining and imaging of control U2OS, and CHCHD4 (WT)-expressing cells demonstrated that the cytoplasmic localisation of vimentin was also regulated by CHCHD4 expression, with a more perinuclear distribution induced by CHCHD4 overexpression (Fig. 5g and Additional File 5: S5e), which has been identified as a consequence of MET [49, 50]. To investigate whether CI activity was required for the CHCHD4-mediated effects on changes in vimentin intracellular distribution, we treated both control U2OS and CHCHD4 (WT)-expressing U2OS cells with rotenone, and assessed vimentin localisation by immunofluorescence. Rotenone treatment led to the redistribution of vimentin from the perinuclear region to the cytoplasm in cells overexpressing CHCHD4 (Fig. 5h), photocopying the vimentin distribution in control U2OS cells. Together, these data show that increased expression of CHCHD4 regulates the EMT phenotype of tumour cells, through its effects on mitochondrial function, and highlights the importance of tumour cell metabolism in the regulation of a proliferative-EMT switch.

## DISCUSSION

Dysregulated metabolism is a common feature of tumour cells, and numerous oncogenes are potent regulators of metabolic pathways that contribute towards tumour cell proliferation. The mechanisms by which tumour cells maintain biosynthesis of macromolecules for division are incompletely understood and are currently the focus of intensive investigation. Here we have identified CHCHD4 as a new regulator of the metabolic drive which supports tumour cell proliferation, in part through its effects on CI-mediated metabolism of glutamine, and consequently the activity of the mTORC1 signalling pathway. Mitochondria promote proliferation in part through ATP synthesis, but tumour cells often adapt their metabolism to decreases their dependence on OXPHOS for ATP homeostasis, and upregulate non-mitochondrial pathways of ATP synthesis, such as glycolysis and anaerobic fermentation of pyruvate [51, 52]. Besides ATP synthesis, mitochondria are also important metabolic hubs for pathways that support biomass accumulation, and these pathways appear to be more acutely sensitive to perturbations in respiratory chain function than ATP synthesis. Indeed, here we found that doses of the CI inhibitor BAY 87 2243 that are insufficient to block OCR were capable of significantly influencing protein translation and tumour cell proliferation. In support of the essential role of mitochondrial metabolism for biomass accumulation, two recent studies have demonstrated that the synthesis of aspartate is an essential function of mitochondrial respiration for proliferating cells [8, 10]. In agreement with these studies, we show here that aspartate supplementation is capable of partially reversing the anti-proliferative effects of both CI inhibition and glutamine withdrawal, and stimulates mTORC1 mediated activation of translation. Since this rescue by aspartate supplementation was only partial, we conclude that other biosynthetic pathways that depend on CI activity and glutamine availability must also contribute to tumour cell proliferation. For example, both NAD and glutamine are directly involved in nucleotide synthesis [18, 53, 54], and nucleotide supplementation in addition to aspartate supplementation may be necessary to fully recapitulate the proliferation of cells treated with CI inhibitors or glutamine withdrawal. For example, pyrimidine synthesis through the uridine salvage pathway relies directly on respiratory chain activity, through the action of the mitochondrial enzyme dihydroorotate dehydrogenase (DHODH) [55]. In cells depleted of mitochondrial DNA (ρ^0^ cells), supplementation with uridine is necessary to rescue the loss of proliferative capacity [56], demonstrating that supporting nucleotide synthesis is an essential role of the mitochondria for proliferating cells. Indeed, transcriptome analysis of patient samples has identified that upregulation of nucleotide biosynthetic genes is one of the most common metabolic alterations across cancer types [57, 58].

Cancer can be considered a disease of two major phenotypes: dysregulated proliferation, and metastatic dissemination of transformed cells to distant sites. Metastasis is usually (but not always) accompanied by the acquisition by tumour cells of traits which decrease cell-cell interactions, and increase motility and resistance to killing by commonly used therapeutics [59]. In recent years, it has become appreciated that mitochondrial-mediated proliferation and EMT phenotypes are inversely related, though regulated by common pathways [38, 39, 41, 42, 60]. For example, a recent analysis of transcriptome data from patient tumour samples identified that gene sets involved in mitochondrial OXPHOS are commonly changed in tumour tissues compared to normal tissue [57]. However, the direction of change was found to be heterogeneous, with upregulation of mitochondrial OXPHOS in 35% of cancer types, and downregulated in 25% of cancer types [57]. Interestingly however, in those cancers with downregulation of mitochondrial OXPHOS, EMT-related genes were the most significantly upregulated cohort, and these changes were correlated with worse outcomes for patients [57]. In agreement with this, we found that *CHCHD4* expression is also positively correlated with mitochondrial OXPHOS and proliferative signalling pathways (e.g. mTORC1, MYC, E2F), while it is negatively correlated with an EMT gene signature in vitro and in patient samples. It would appear then that by influencing proliferation which depends on mitochondrial activity, CHCHD4 also regulates the EMT phenotype of tumour cells through transcriptional and post-transcriptional mechanisms (e.g. vimentin intracellular distribution).

Our results suggest a paradox with respect to the importance of CHCHD4 in cancer. While we show here that increased *CHCHD4* expression correlates with decreased EMT gene expression, we have already demonstrated that above median *CHCHD4* expression is correlated with increased tumour grade and decreased survival [28]. Since metastatic disease is a common feature of disease progression [59], how then can elevated *CHCHD4* be associated with worse outcomes for patients in certain cancers? The answer may come from our previous work which identified CHCHD4 as a critical regulator of HIF-mediated transcriptional responses to hypoxia [28, 29]. The tumour microenvironment appears to be the primary driver of EMT, since no recurrent mutations in EMT-regulating genes have been identified from genomic sequencing of tumour cells, unlike the myriad mutations in oncogenes and tumour suppressors which regulate proliferation [59]. One significant environmental stimulus of EMT is hypoxia, which is a common feature of tumour tissues which outgrow their vascular supply [61, 62]. Activation of HIF-signalling under hypoxia leads to transcriptional activation of EMT-related genes such as vimentin and N-cadherin, along with the suppression of MET-related genes such as E-cadherin [63]. Furthermore, HIF-signalling decreases mitochondrial OXPHOS by diverting carbon fuels away from metabolism by the mitochondria [51, 64], and in some cases decreases mitochondrial mass by suppressing mitochondrial biogenesis [65]. Thus, in tumours with elevated CHCHD4 expression, the increased proliferation it affords may increase the size of the hypoxic niche, while simultaneously enhancing HIF activation, and consequently metastatic dissemination of tumour cells. Indeed, we have shown that silencing of *CHCHD4* significantly decreases the hapto-migration and invasion of HCT116 cells in hypoxia [28], demonstrating the relationship between CHCHD4 and EMT phenotypes in hypoxia. It will be important to make more detailed investigations into the influence of CHCHD4 on hypoxia responses and metastatic phenotypes in 3D culture models and patient tumours, to fully understand the contribution of CHCHD4 to disease progression.

From a therapeutic perspective, mitochondrial metabolism is an attractive target for treatment regimes, but is not without significant risk of toxicity. Agents which target the mitochondria are potently anti-proliferative, and several small molecules are in clinical trials which inhibit mitochondrial metabolic pathways [14, 15, 23]. The present study points to a potential complication with this kind of therapeutic strategy however; in that targeting mitochondrial metabolism (e.g. at CI) may stimulate tumour cells to develop EMT characteristics which promote metastatic dissemination. Primary tumour cells which evade killing by chemotherapies may therefore contribute to the metastatic population of transformed cells, and anti-proliferative agents may in fact contribute to disease progression and relapse. It will be important to thoroughly understand the molecular mechanism which underpin this apparent inverse relationship between proliferation and EMT in order to be able to devise treatment regimens which effectively and systemically remove transformed cells from patients.

## CONCLUSIONS

Mitochondrial metabolism plays a central role in tumour cell proliferation. Our current study demonstrates that the mitochondrial import protein CHCHD4 regulates tumour cell proliferation through its effects on CI expression and activity. Increased CI activity increases metabolism of glutamine, and drives mTORC1-mediatedsignalling. Furthermore, our study demonstrates that CHCHD4 expression regulates the EMT phenotype of tumour cells, and is negatively correlated with the expression of EMT genes in normoxia. Future studies will further investigate the influence of CHCHD4 on the metabolic landscape of cultured tumour cell lines (e.g. nucleotide synthesis), and the proliferation and metastatic behaviour of tumour cells in vitro and in vivo.

## METHODS

### Cell culture

Human U2OS, HeLa, MCF7, HCT116, U87-MG and PC3 cell lines were all obtained from American Tissue Culture Collection (ATCC). Human osteosarcoma U2OS control and independent clonal cell lines (WT.cl1 and WT.cl3) expressing CHCHD4.1 cDNA (CHCHD4-WT-expressing cells) or CHCHD4-C66A/C668A cDNA (CHCHD4-(C66A/C68A)-expressing cells) have been described by us recently [29]. Human HCT116 cells were used to generate a stable clonal cell line (WT.cl8)-expressing CHCHD4.1 cDNA (CHCHD4 (WT)-expressing cells) cDNA by puromycin selection using constructs we have described previously [28]. Human U2OS or human HCT116 colon carcinoma cells were used to stably express empty GFP Vector (shRNA control 2)) or two independent shRNAs targeting CHCHD4 (CHCHD4 shRNA1 or CHCHD4 shRNA2) utilizing a green fluorescent protein (GFP)-SMARTvector^TM^ pre-packaged lentivirus system from ThermoFisher Scientific. Independent cell pools were selected, expanded and characterized. All cell lines were maintained in Dulbecco’s modified eagle medium (DMEM) containing glucose (4.5 g/L) (Life Technologies), and supplemented with 10% fetal calf serum (FCS, SeraLabs), penicillin (100 IU/mL), streptomycin (100 μg/mL) and glutamine (6 mM), all purchased from Life Technologies. Cell lines used were authenticated and routinely confirmed to be negative for any mycoplasma contamination.

### Antibodies and Reagents

The rabbit polyclonal P-p70S6K (#9205), T-p70S6K (##2708), LC3B (#3868), SDHA (#11998), COXIV (#4850), MYC-tag (2272), Vimentin (#3932), and N-cadherin (#13116) antibodies were purchased from Cell Signaling Technology. The rabbit polyclonal CHCHD4 (HPA034688) antibody was purchased from Cambridge Biosciences. The rabbit polyclonal NDUFB10 (ab196019), NDUFS3 (ab110246), UQCRC2 (ab14745) and mouse monoclonal α-Tubulin (ab7291), and β-actin (ab6276) antibodies were purchased from Abcam. The goat anti-mouse IgG Alexa Fluor 568 (A11031, 1:1000) and goat anti-rabbit IgG Alexa Fluor 488 (A11034, 1:1000) were purchased from ThermoFisher Scientific. The rabbit anti-puromycin antibody was a gift from Stefan Marciniak (CIMR, Cambridge). Uniformly labelled ^13^C5-glutamine (CLM-1822-H-MPT-PK) purchased from CK Isotopes. Rotenone, antimycin A, DAPI nitrotetrazolium blue, NADH, aspartate, L-lysine, L-arginine, L-lysine-^13^C_6_, ^15^N_2_ and L-arginine-^13^C_6_, ^15^N_4_ (Arg-10) were purchased from Sigma Aldrich. BAY 87 2243 was purchased from MedChemExpress.

### Gene expression analysis

Total RNA samples were isolated using the GeneElute kit, following the manufacturer’s protocol (Sigma-Aldrich). cDNA synthesis was carried out using the qScript synthesis kit, following the manufacturer’s protocol (Quantabio). mRNA expression was measured by quantitative (Q)-PCR using SYBR Green Mastermix (Eurogentec Ltd.) and the DNA Engine Opticon 2 system (BioRad). Q-PCR primer sequences as follows: *CHCHD4*_F 5’-GAGCTGAGGAAGGGAAGGAT-3’; *CHCHD4*_R 5’-AATCCATGCTCCTCGTATGG-3’; *KRT18*_F 5’-TAGATGCCCCCAAATCTCAG-3’; *KRT18*_R 5’-CACTGTGGTGCTCTCCTCAA-3’; *CDH2*_F 5’-AGGATCAACCCCATACACCA-3’; *CDH2*_R 5’-TGGTTTGACCACGGTGACTA-3’; *VIM*_F 5’-GAGAACTTTGCCGTTGAAGC-3’; *VIM*_R 5’-TCCAGCAGCTTCCTGTAGGT-3’; *ACTB*_F 5’-CCCAGAGCAAGAGAGG-3’; *ACTB*_R 5’-GTCCAGACGCAGGATG-3’.

### SILAC

Cells were incubated in arginine and lysine free DMEM (Life Technologies), supplemented with either L-lysine and L-arginine (light) or L-lysine-^13^C_6_, ^15^N_2_ (Lys-8) and L-arginine-^13^C_6_, ^15^N_4_ (Arg-10) (heavy) stable isotope labelled amino acids (Sigma Aldrich). Media was also supplemented with 10% dialysed FCS (Sigma), penicillin (100 IU/mL), streptomycin (100 μg/mL) and L-glutamine (200 mM), and all purchased from Life Technologies. Amino acid incorporation was carried out over >5 passages. Mitochondrial fractions were isolated as described below. 50 μg of enriched mitochondria were resolved approximately 6 cm into a pre-cast 4–12% Bis-Tris polyacrylamide gel (ThermoFisher Scientific). The lane was excised and cut in 8 approximately equal chunks and the proteins reduced, alkylated and digested in-gel. The resulting tryptic peptides were analysed by LC-MSMS using a Q Exactive coupled to an RSLCnano3000 (ThermoFisher Scientific). Raw files were processed using MaxQuant 1.5.2.8 using Andromeda to search a human Uniprot database (downloaded 03/12/14). Acetyl (protein N-terminus), oxidation (M) and deamidation (N/Q) were set as variable modifications and carbamidomethyl (C) as a fixed modification. SILAC data was loaded in R to process it with the Microarray-oriented limma package to call for differential expression [66], relying on the original normalisation processes produced by MaxQuant as reported previously [67, 68]. Three independent SILAC experiments using control U2OS and CHCHD4 (WT.cl1)-expressing cells involving parallel labelling were performed. Two independent SILAC experiments using shRNA control and CHCHD4 (shRNA) knockdown cells involving parallel labelling were performed.

### Sulforhodamine B (SRB) viability assay

Cells were plated in appropriate tissue culture vessels, and allowed to adhere overnight prior to treatment. At the end of incubation, media was removed and cells were fixed with 10% trichloroacetic acid (TCA) for 30 min. TCA was washed with water, wells were allowed to air dry, and then an excess of 0.4% (w/v) SRB in 1% acetic acid was used to stain fixed cells for >10 min. Excess SRB was washed off with 1% acetic acid solution. Bound SRB was resuspended in a suitable volume of 10 mM Tris, and absorbance of solution measured at 570 nm. For proliferation assays, cells were plated on ‘day −1’ in triplicate in 12 well plates, and cultured in maintenance DMEM overnight, after which ‘day 0’ plates fixed with TCA. For drug sensitivity assays, cells were plated in triplicate columns in 96-well plates, and cultured in maintenance DMEM overnight. Appropriate wells were dosed with serial dilutions of compounds, including vehicle control wells. Cells were incubated for desired time points followed by SRB assay. GI50 values were calculated using regression analysis

### BN-PAGE and CI assay

Samples were prepared for BN-PAGE using the Native-PAGE Sample Preparation Kit and Protocol (Life Technologies) using a 10% dodecylmaltoside (DDM) permeabilization solution. Samples were run on 3-12% gradient non-reducing acrylamide gels (Life Technologies). For complex I activity assay, samples were run without coomassie blue, and gels incubated in a complex I assay buffer (100 μM NADH and 0.5 mg/mL nitrotetrazolium blue in 20 mM Tris) as previously described [69].

### Microscopy

For immunofluorescence microscopy, cells were seeded onto 13mm diameter coverslips, and after treatment, were fixed in ice-cold methanol overnight at −20°C. Coverslips were then washed with phosphate buffered saline (PBS). Immunostaining was carried out by serial incubation using primary antibodies directed to Vimentin (rabbit polyclonal) and α-Tubulin (mouse monoclonal), followed by an anti-rabbit Alexa 488 and anti-mouse Alexa-568 secondary antibodies, as well as DAPI (1 μg/mL). All cell imaging was carried out using a DMI4000 B inverted microscope (Leica). Vimentin distribution analysis was carried out using Cell Profiler Image analysis software as previously described [29]. For live cell imaging, cells were imaged using a Nikon

### Respirometry

Oxygen consumption rates (OCR) were determined using a Seahorse XF96 Analyser (Seahorse Bioscience). Respiratory profiles were generated by serial treatment with optimised concentrations of oligomycin (1 µg/mL), p-[trifluoromethoxy]-phenyl-hydrazone (FCCP, 500 nM), and rotenone (500 nM). Cell number normalisation was carried out post-respirometry using sulforhodamine B (SRB) staining of TCA fixed cells in the assay plate.

### Gene Set Analysis

Transcriptomic data generated by TCGA was accessed from the CBioPortal data portal (http://www.cbioportal.org/). Samples for gene expression heatmaps were filtered by excluding samples with a z-score value for *CHCHD4* expression >1.5. Genes for correlation analyses were filtered by excluding samples with a *p*-value >0.05. All correlations were calculated using Spearman’s method. Gene set enrichment analyses were carried out at the Broad Institute analysis portal (http://software.broadinstitute.org/gsea/msigdb/annotate.jsp).

### Western blot densitometry

Western blot signal intensity was measured per lane using ImageJ (NIH) analysis software. Phosphorylated (P)-p70S6K band intensities were normalised to total (T)-p70S6K band intensities, then relative band intensities were calculated compared to untreated samples for each cell line.

### Metabolomics Analysis

For steady-state metabolomics or metabolite tracing experiments 1 × 10^5^ cells were seeded in 6 wells of a 6-well plate for each cell line. After 24h cells were washed twice with PBS and medium was changed with DMEM medium containing metabolite tracers. For glutamine tracing experiments 6 mM ^13^C5-glutamine was added to glutamine-free DMEM, together with 10% v/v FBS. After incubation for 24h with medium containing metabolite tracers, one well was used to estimate cell number. To extract intracellular metabolites, cell plates were placed on ice, washed twice with ice-cold PBS and 1 mL of MEB per 10^6^ cells was added to each well and cells were scraped. One cycle of freeze-thawing at −80°C was performed to further lyse the cells. Intracellular fractions were then incubated in a thermomixer (Eppendorf) at max speed for 15 minutes at 4°C. Proteins were then pelleted by centrifuging samples at 16000 g for 10 min at 4°C and supernatants were transferred into glass vials and stored at −80°C until further analysis. Liquid chromatography–mass spectrometry (LCMS) analysis was performed on a QExactive Orbitrap mass spectrometer coupled to a Dionex UltiMate 3000 Rapid Separation LC system (Thermo). The LC system was fitted with a SeQuant ZIC-pHILIC (150mm × 2.1mm, 5mm) with the corresponding guard column (20m × 2.1mm, 5mm) both from Merck. The mobile phase was composed of 20cmM ammonium carbonate and 0.1% ammonium hydroxide in water (solvent A), and acetonitrile (solvent B). The flow rate was set at 200 mL/min with a previously described gradient [70]. The mass spectrometer was operated in full MS and polarity switching mode scanning a range of 50-750 m/z. Samples were randomized, in order to avoid machine drift, and were blinded to the operator. The acquired spectra were analyzed using XCalibur Qual Browser and XCalibur Quan Browser software (Thermo Scientific) by referencing to an internal library of compounds. Calibration curves were generated using synthetic standards of the indicated metabolites. Intensity of intracellular metabolites were normalized on total ion sum (normalized intensity values). For interpretation of labeling patterns normalized intensities of isotopologues were further normalized on total isotopologue sum for each metabolite species (proportion of total pool values).

## DECLARATIONS

### Availability of data and materials

Requests can be made to the corresponding author relating to materials generated in this study.

### Authors’ contributions

LWT designed and performed experiments, analysed data and wrote the manuscript. JS and CE designed and performed experiments, analysed data. ASHC and CF performed metabolomics experiments. TSB and GS assisted with NAD/NADH imaging analysis. MA provided the concept for the study, designed experiments, analysed data, wrote the manuscript and acquired funding. All authors reviewed and edited the manuscript.

### Ethics approval

Not applicable

