## Supplemental Figure 1 for "CHCHD4 regulates a proliferation-EMT switch in tumour cells, through respiratory complex I mediated metabolism"

a

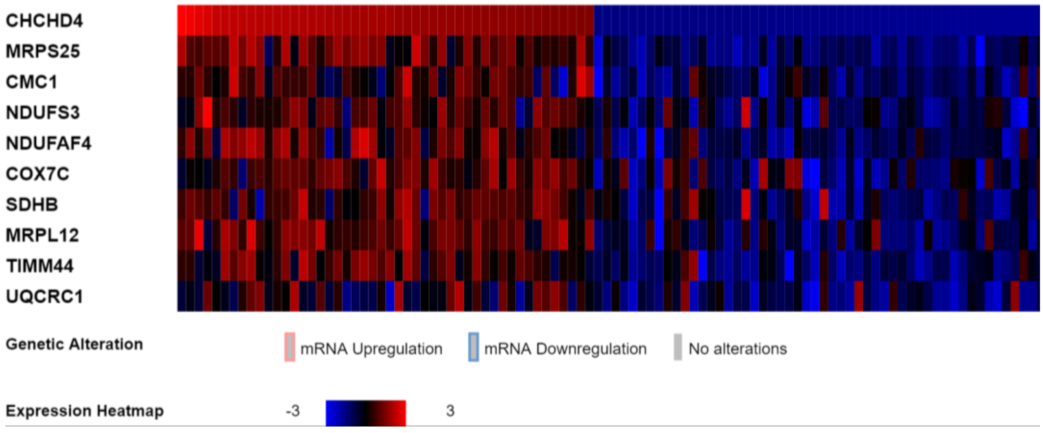


b

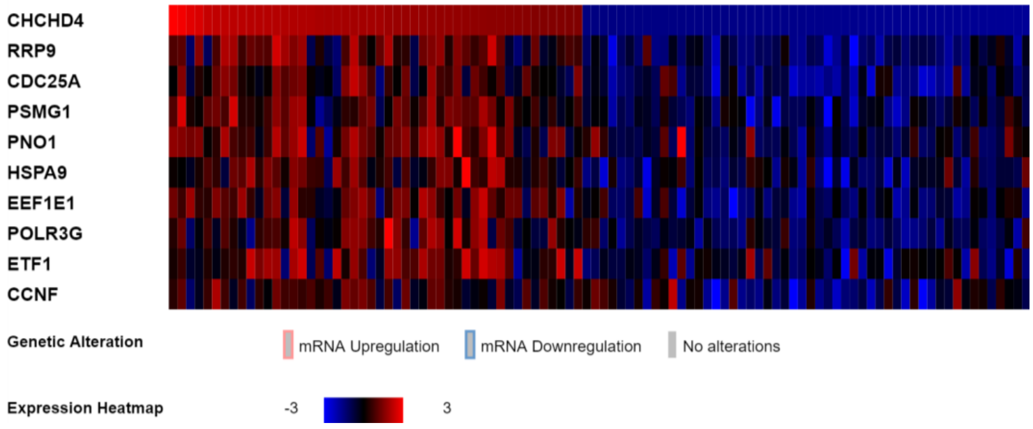


c

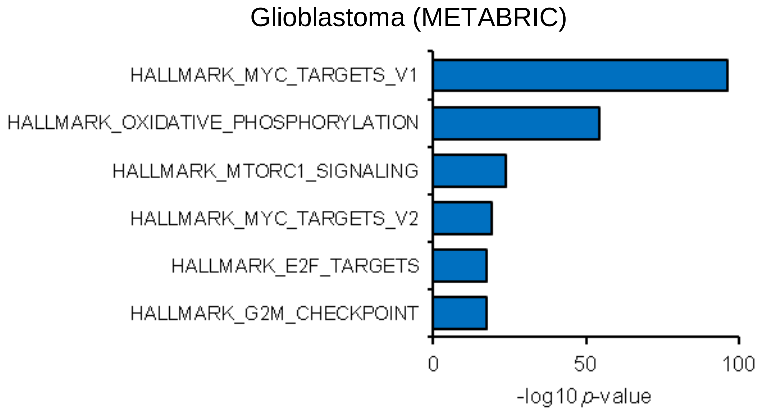


**Figure S1. CHCHD4 expression is associated with proliferative signalling in tumours.**

**Figure S1. CHCHD4 expression correlates with mitochondrial OXPHOS and proliferative gene signatures in vitro and in vivo. a** Heatmap of selected genes from HALLMARK_OXIDATIVE_PHOSPHORYLATION gene-set (Broad Institute) that are positively correlated with CHCHD4 expression in Novartis/Broad Institute Cancer Cell Line Encyclopedia RNASeq data. n=967 cell lines. **b** Heatmap of selected genes from HALLMARK_MTORC1_SIGNALLING gene set (Broad Institute) that are positively correlated with CHCHD4 expression in Novartis/Broad Institute Cancer Cell Line Encyclopedia RNASeq data. n=967 cell lines. **c** Chart shows GSEA of genes positively correlated with CHCHD4 expression in glioblastoma patient tumours.
