## Supplemental Figure 2 for "CHCHD4 regulates a proliferation-EMT switch in tumour cells, through respiratory complex I mediated metabolism"

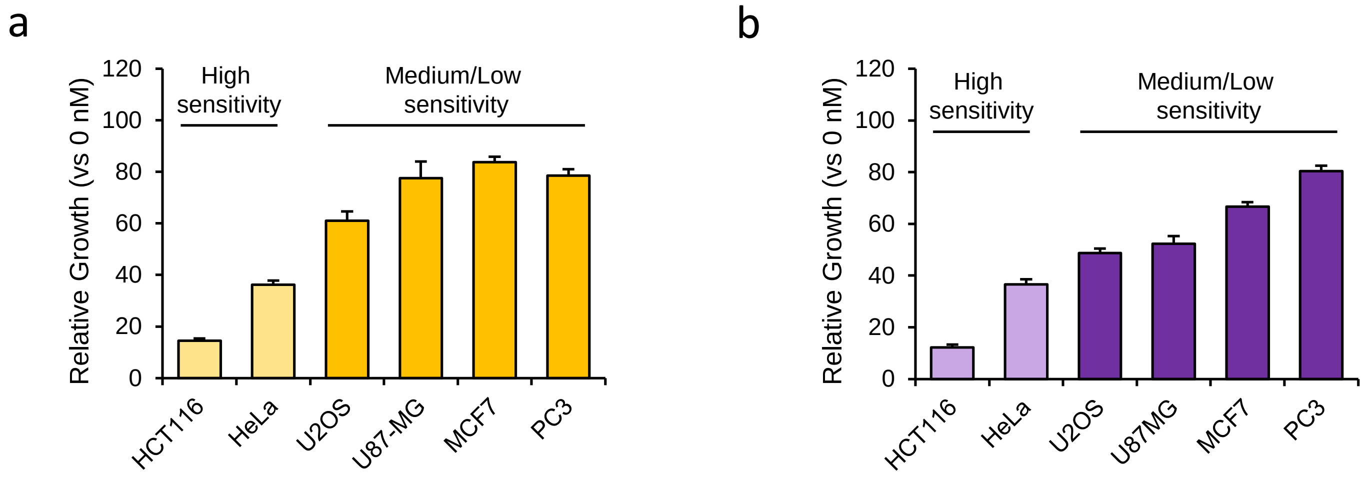


**Figure S2. Tumour cell proliferation is correlated with CHCHD4 expression and mitochondrial function.**

**Figure S2. Tumour cell proliferation correlates with CHCHD4 expression and mitochondrial function. a** Chart shows growth of tumour cell line panel treated with 500 nM BAY 87 2243 for 72h, relative to untreated (0 nM) cells. ±SD. n=3. **b** Chart shows growth of tumour cell line panel treated with 3 µM antimycin A for 72h, relative to untreated (0 µM) cells. ±SD. n=3.
