## Supplemental Figure 3 for "CHCHD4 regulates a proliferation-EMT switch in tumour cells, through respiratory complex I mediated metabolism"

a

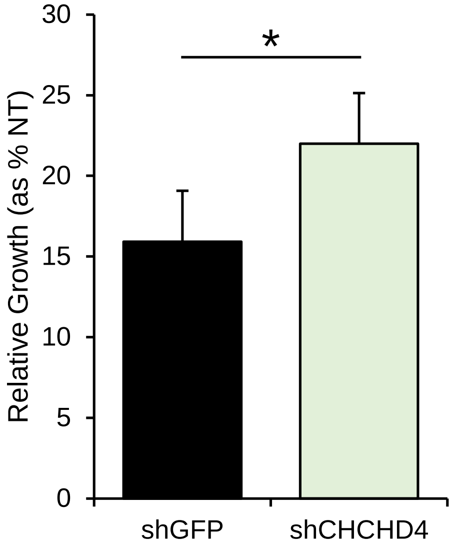


**Figure S3. CHCHD4 regulates CI expression and activity in tumour cells, and regulates the mTORC1 pathway.**

**Figure S3. CHCHD4 regulates CI expression and activity in tumour cells, and regulates the mTORC1 pathway. a** Chart shows growth of HCT116 cells stably expressing control shRNA (shGFP) or CHCHD4-targeting shRNA (shCHCHD4) treated with 100 nM rotenone for 72h relative to untreated cells (NT). ±SD. n=3.
