## Supplemental Figure 4 for "CHCHD4 regulates a proliferation-EMT switch in tumour cells, through respiratory complex I mediated metabolism"

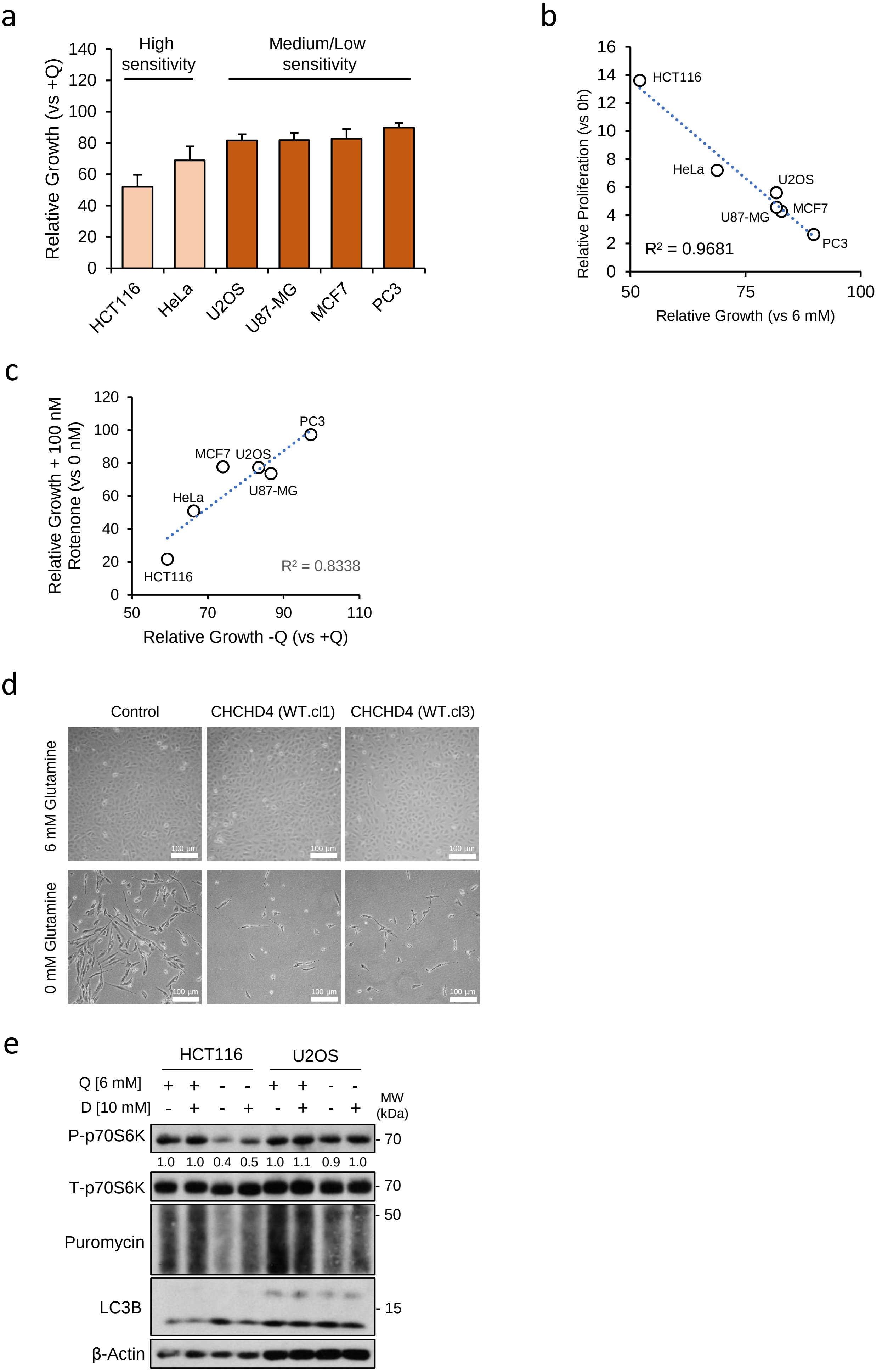


**Figure S4. CHCHD4 regulates tumour cell proliferation and glutamine consumption.**

**Figure S4. CHCHD4 regulates proliferation and glutamine consumption. a** Chart shows growth of tumour cell line panel incubated in the absence of glutamine (Q) for 72h, relative to incubation with 6 mM glutamine (+Q). ±SD. n=3. **b** Chart shows xy scatter of proliferation rates of indicated cell lines at 72h, and growth rates of cells incubated in the absence of glutamine relative to incubation with 6 mM glutamine. Trend line (dashed blue) and R2 value (Spearman's correlation) shown. **c** Chart shows xy scatter of growth rates of indicated cell lines treated with 100 nM rotenone relative to untreated (0 nM), and in the absence of glutamine (-Q) relative to incubation with 6 mM glutamine (+Q). Trend line (dashed blue) and R2 value (Spearman's correlation) shown. **d** Bright field microscopy images of control U2OS cells and cells overexpressing wild-type CHCHD4 (WT.cl1, WT.cl3) after 10 days cultured with 6 mM and 0 mM glutamine. **e** Western blot shows levels of phosphorylated (P-) and total (T-) p70S6K, puromycin labelled proteins, and LC3B in HCT116 and U2OS cells treated as indicated for 24h. β-Actin used as load control. Relative band intensities of P-p70S6K indicated.
