## Supplemental Figure 5 for "CHCHD4 regulates a proliferation-EMT switch in tumour cells, through respiratory complex I mediated metabolism"

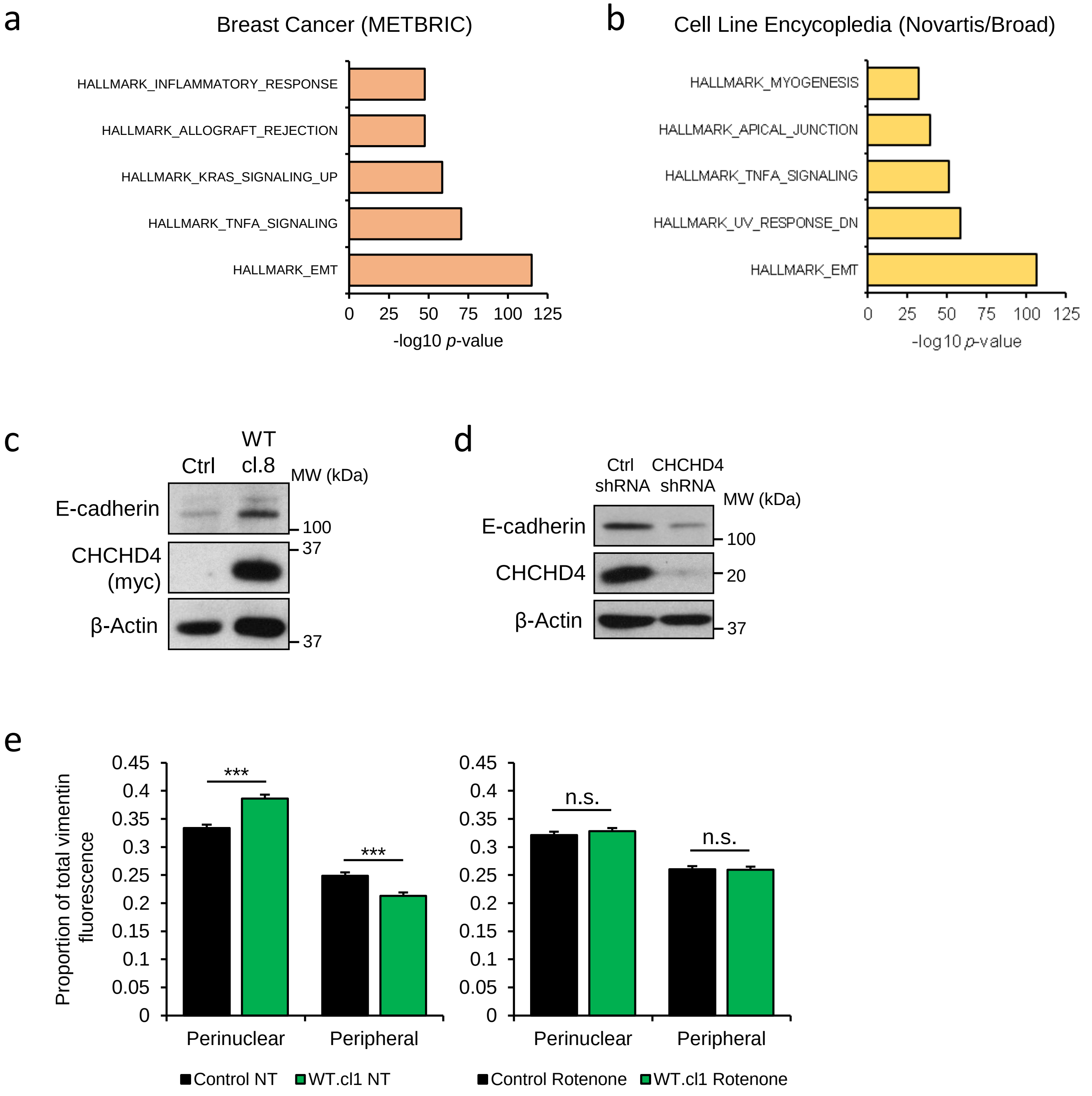


**Figure S5. CHCHD4 regulates the EMT phenotype of tumour cells.**

**Figure S5. CHCHD4 regulates the EMT phenotype of tumour cells. a** Chart shows GSEA of genes negatively correlated with CHCHD4 expression in breast cancer patient tumours. **b** Chart shows GSEA of genes negatively correlated with CHCHD4 expression in Novartis/Broad Institute Cell Line Encyclopedia. n=967 cell lines. **c** Western blot shows levels of E-cadherin and myc-tagged CHCHD4 in control (Ctrl) HCT116 cells, and cells overexpressing wild-type CHCHD4 (WT.cl8). β-Actin used as load control. **d** Western blot shows levels of E-cadherin and CHCHD4 in HCT116 cells expressing control (Ctrl) shRNA or CHCHD4-targeting shRNA. β-Actin used as load control. **e** Chart shows relative proportion of fluorescently labelled vimentin in the perinuclear and peripheral sections of control U2OS cells and cells overexpressing wild-type CHCHD4 (WT.cl1) untreated (NT) or treated with 50 nM rotenone for 72h. ±SD. n=2 incubations, 5 fields of view per condition.
